# Transcriptome Analysis of Adult *C. elegans* Cells Reveals Tissue-specific Gene and Isoform Expression

**DOI:** 10.1101/232728

**Authors:** Rachel Kaletsky, Vicky Yao, April Williams, Alexi M. Runnels, Sean B. King, Alicja Tadych, Shiyi Zhou, Olga G. Troyanskaya, Coleen T. Murphy

**Affiliations:** Department of Molecular Biology, Princeton University, Princeton NJ 08544; Department of Computer Science, Princeton University, Princeton, NJ 08544, USA; Flatiron Institute, Simons Foundation, New York, NY 10010, USA; Lewis-Sigler Institute for Integrative Genomics, Princeton University, Princeton, NJ 08544, USA

**Author notes:** Equal contribution (Rachel Kaletsky, Vicky Yao).

**Keywords:** cell-specific transcription, tissues, RNA-seq, adult, *C. elegans*, human, IIS, Insulin, DAF-16, FOXO, CREB, TGF-b

## Abstract

The biology and behavior of adults differ substantially from those of developing animals, and cell-specific information is critical for deciphering the biology of multicellular animals. Thus, adult tissue-specific transcriptomic data are critical for understanding molecular mechanisms that control their phenotypes. We used adult cell-specific isolation to identify the transcriptomes of *C. elegans’* four major tissues (or “tissue-ome”), identifying ubiquitously expressed and tissue-specific “super-enriched” genes. These data newly reveal the hypodermis’ metabolic character, suggest potential worm-human tissue orthologies, and identify tissue-specific changes in the Insulin/IGF-1 signaling pathway. Tissue-specific alternative splicing analysis identified a large set of collagen isoforms and a neuron-specific CREB isoform. Finally, we developed a machine learning-based prediction tool for 70 sub-tissue cell types, which we used to predict cellular expression differences in IIS/FOXO signaling, stage-specific TGF-b activity, and basal vs. memory-induced CREB transcription. Together, these data provide a rich resource for understanding the biology governing multicellular adult animals

## Introduction

Animals progress through many stages of development before reaching adulthood, and as adults, they exhibit metabolic and behavioral differences from still-developing animals. Studies in the nematode *C. elegans* demonstrate this phenomenon well: both biological responses and gene expression differ significantly in different stages (Spencer et al., 2011; Kaletsky et al., 2015). Therefore, to understand the biology underlying tissue-specific adult behavior, it is critical to identify adult, tissue-specific transcriptomes.

The advent of whole-genome gene expression approaches allowed the identification of a cell’s full set of mRNA transcripts, ushering in a new era of understanding biological dynamics (DeRisi et al., 1997). The ongoing development of new methods to isolate and sequence individual cells in order to approximate their metabolic and biochemical state has refined our understanding of single cells (Shalek et al., 2013). The next frontier in this work is the gene expression analysis of whole animals on a tissue-by-tissue and cell-by-cell basis.

*C. elegans* is the simplest multicellular model system, with only 959 somatic (non-germline) cells in the fully developed adult animal. Four tissues—muscles, neurons, intestine, and the epidermis (or “hypodermis”)—comprise the bulk of the animal’s somatic cells and are largely responsible for the animal’s cell autonomous and non-autonomous biological regulation. Until recently, most transcriptional analyses of *C. elegans* adults utilized whole worms, but the need to identify tissue-specific transcripts in order to better understand both tissue-specific and non-autonomous signaling has become apparent. Tools used to isolate embryonic and larval stage *C. elegans* cells (Cao et al., 2017; Fox et al., 2005; Spencer et al., 2011, 2014; Zhang et al., 2002) have allowed the transcriptional profiling of specific tissues and cell types, shedding light on larval development processes, but lack information specific to adult tissues. Much of worm behavioral analysis, and all aging studies—for which *C. elegans* is a premier model system (Kenyon, 2010)—is performed in adults, which are less amenable to standard isolation approaches due to their tough outer cuticle. Therefore, we developed a method to disrupt and isolate adult tissues (Kaletsky et al., 2015). That work revealed that the adult neuronal transcriptome differs significantly from earlier embryonic and larval stages, and that the adult neuronal transcriptome best reveals genes involved in behavior and neuronal function. The other major tissues—muscle, intestine, and hypodermis—are likely to provide insight into important adult-specific processes that are widely studied in *C. elegans* as models of human biology, such as pathogenesis, reproduction, age-related decline, and others.

Here we have performed cell-specific transcriptional analysis on the four major somatic tissues isolated from adult worms, and characterized the tissues using these data. We used the set of highly enriched tissue gene sets to identify tissue-specific DAF-16 transcriptional targets and to identify transcriptional parallels between worm and human tissues. Additionally, our sequencing method allowed the identification of tissue-specific alternatively spliced gene isoforms. Finally, we present a method to predict gene expression in 70 different subtissue cell types, and demonstrate its utility in the characterization of potential cellular differences in gene expression for several different signaling pathways. Together, these data provide a rich resource for the examination of adult gene expression in *C. elegans*.

## Results

### Isolation and sequencing of major adult *C. elegans* tissues

To identify the transcriptomes of adult *C. elegans* tissues, it is necessary to break open the outer cuticle and release, filter, and sort cells while minimizing cell damage (Kaletsky et al., 2015). Day 1 adult animals with fluorescently-marked neurons (*Punc-119::gfp*), muscle (*Pmyo-3::mCherry*), hypodermis (*pY37A1B.5::gfp*), and intestine (*Pges-1::gfp;* Figure 1A) were briefly subjected to SDS-DTT treatment, proteolysis, mechanical disruption, cell filtering, FACS, RNA amplification, library preparation, and single-end (140 nt) Illumina sequencing, as previously described (Kaletsky et al., 2015). We collected 27 adult tissue samples (7 neuron, 5 intestine, 7 hypodermis, 8 muscle). For each sample, approximately 50,000 to 250,000 GFP or mCherry positive events were collected, yielding ~5 to 25 ng total RNA. 35 – 200 million reads (average of 107,674,388 reads) were obtained for each sample, and mapped to the *C. elegans* genome (Ensembl release 84).

**Figure 1:**
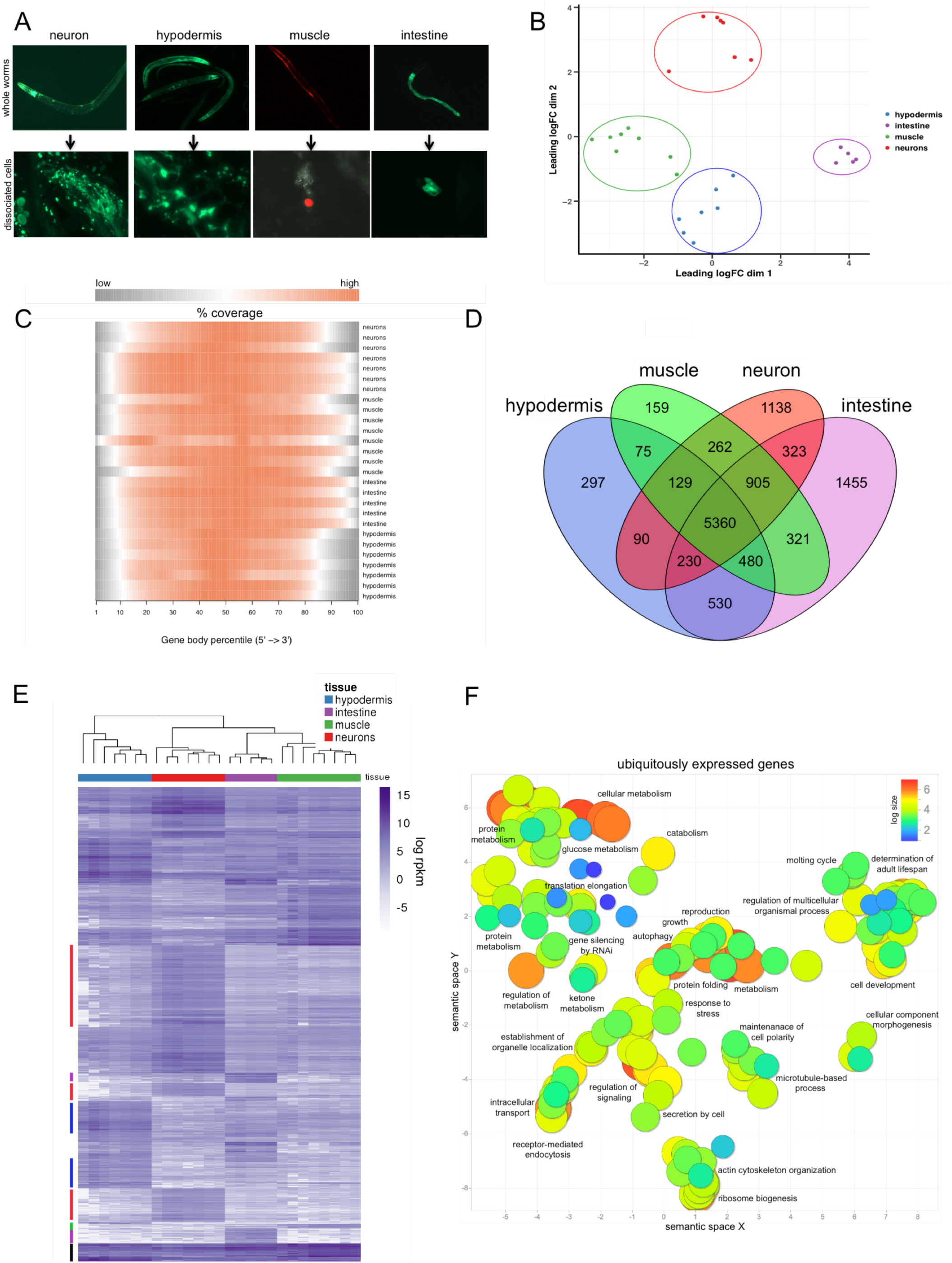
Sorting and RNA-seq of isolated adult *C. elegans* tissues. A) Day 1 adult *C. elegans* strains used in our study to sort and identify neurons (*Punc119::GFP*), hypodermis (*pY37A1B.5::GFP*), muscle (*Pmyo-3::mCherry*), and intestine (*Pges-1::GFP*) before (top) and after cell disruption (bottom, prior to debris filtering). B) Multidimensional scale plot of all samples used in this study. C) Heatmap showing read coverage profiles over gene body to evaluate whether coverage is uniform (versus potential 5’ or 3’ bias). D) Venn comparison of expressed genes in each tissue. E) Heatmap of gene expression across tissues. Log rpkm expression values for the top 2000 differentially expressed genes per tissue (blue = high, white = low). Color bars (left) represent clusters of tissue specific gene expression (ubiquitous, black; hypodermis, blue; neuron, red; intestine, purple; muscle, green). F) Gene ontology (GO) analysis of ubiquitously expressed genes (defined as the 5,360 genes (see Venn above) expressed across all 4 tissues. GO terms with padj < 0.05 (hypergeometric test) were plotted using REVIGO (Supek et al., 2011).

Multidimensional scaling analysis (Figure 1B) that the samples cluster best with their respective tissue types, and that muscle and hypodermis are most closely related, while neuronal and intestine samples are more distinct from one another. Subsampling analysis (Robinson and Storey, 2014), which determines whether sequencing has been performed to sufficient depth, suggests that this estimate of gene expression is stable across multiple sequencing depths (Figure S1A), and thus gene expression differences represent true differences between tissues.

We obtained reads across the whole transcript length (rather than selecting the 3’ end of mRNA via the polyA tail) in order to analyze tissue-specific isoform expression (see below). To assess RNA degradation in each sample, we determined the gene body coverage for all 20,389 protein-coding genes (Wang et al., 2012); the transcripts have consistent, uniform coverage, with best coverage within the gene bodies (Figure 1C).

“Expressed” genes are defined as those with both (1) an average log(rpkm) greater than 2, and (2) with each replicate of that tissue having a log(rpkm) greater than 1, resulting in the detection of 8437 neuron, 7691 muscle, 7191 hypodermis, and 9604 intestine protein-coding genes (Figure 1D, Table S1); 5360 genes are expressed in all sampled tissues. Hierarchical clustering of the top 2000 differentially-expressed genes per sample across the four tissue types shows that intra-group tissue samples are most similar, specific genes characterize particular tissue types (especially neurons), and that there is a subgroup of genes expressed in all tissues (Figure 1E). As expected, Gene Ontology (GO) analysis of the ubiquitously-expressed gene set shows that basic cell biological and metabolic processes are shared, including such terms as *intracellular transport*, *protein metabolism*, *catabolism*, *glucose metabolism*, *ribosome biogenesis*, *translation elongation*, *maintenance of cell polarity*, and *microtubule-based process* (Figure 1F; Table S2). Additionally, terms associated with protection of the cell, such as *response to stress*, *autophagy*, *protein folding*, *gene silencing by RNAi*, and *determination of adult lifespan* appear in the ubiquitous category.

### Adult tissues express a set of genes distinct from larval tissues

We previously observed that embryonic and larval transcriptomes lack information relevant to adult neuronal activity and behaviors (Kaletsky et al., 2015). The addition of four neuronal samples here did not assessment; more than four thousand transcripts were found in adult neurons that were not found in embryonic or larval analyses (Figure 2A), and those GO terms are related to neuronal function, while embryonic and larval genes (Spencer et al., 2011) are associated with development (Kaletsky et al., 2015) (Table S3). Similarly, the hypodermis, muscle, and intestine all express transcripts that were not significantly detected in earlier life stages, including GO terms associated with *determination of adult lifespan* and other adult functions (Figure 2A, Tables S4-S6). Furthermore, in each adult tissue we see examples of highly expressed genes that have known associations with adult behaviors and functions, including reproduction, germline maintenance, mating, egg laying, and longevity. For example, adult-enriched genes include neuronal expression of neuropeptides (*nlp-18, -11, -14, -21*), FMRF-like peptides (*flp-1, -9 -19, -28, -3*); synaptobrevin (*snb-1*), the sodium/potassium ATPase EAT-6, which regulates feeding, fertility, and lifespan (Lakowski and Hekimi, 1998), and *pdf-1*, which is involved in mate searching (Barrios et al., 2012). The adult hypodermis expresses higher levels of lifespan regulators (*hsp-1*, *sip-1*, *dod-6*, and the 14-3-3 protein *ftt-2*), as well as metabolic enzymes (*mdh-1* and *-2*, *idh-1*, *enol-1*, *qdpr-1*, *tdo-2*). Intestinal adult targets include *abu-11*, which is induced by ER stress and whose overexpression increases lifespan (Viswanathan et al., 2005), and *ril-1*, which regulates lifespan (Hansen et al., 2005). Several adult tissues express higher levels of secreted fatty acid and retinol-binding proteins (*far*- genes), specific ribosomal subunits, and cytoskeletal elements (e.g., *unc-54*, *tni-3*, *and act-1, -2*).

**Figure 2:**
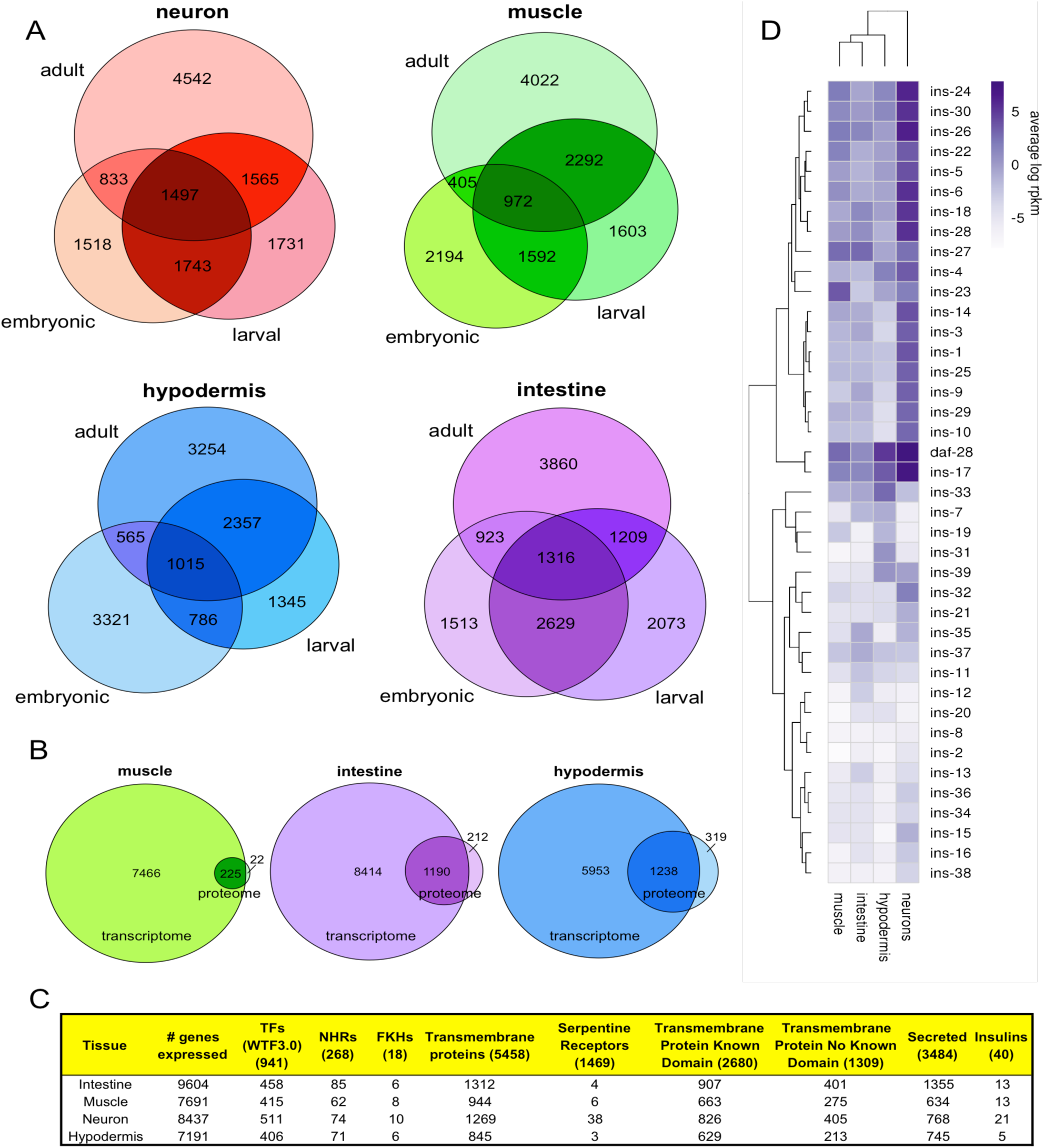
Tissue specificity of gene expression. A) Comparison of expressed genes in larval, embryonic (Spencer et al., 2011), and adult for each tissue (neuron, muscle, hypodermis, intestine) with expression cutoffs as described in Spencer et al., 2011. B) Comparison of expressed genes with proteomic data in three adult tissues (Reinke et al., 2017). Proteins were classified as “expressed” if present in all (3 out of 3) biological replicates. Cytosolic and nuclear proteins were pooled for this comparison. C) Classes of genes expressed in each tissue type; original gene category lists from (Pierce et al., 2001; Reece-Hoyes et al., 2005; Suh and Hutter, 2012). TF, transcription factors; NHR, nuclear hormone receptors; FKH, forkhead transcription factor family. D) Heatmap of insulin expression across adult tissues (average log rpkm; blue = high; white = low).

### Tissue-specific gene and protein expression largely overlap

Recently, Reinke, et al. (2017) used a proximity labeling-based proteomics method to identify ~3000 proteins in four large non-neuronal tissues of the worm, three of which (muscle, intestine, and hypodermis) are shared with our analysis. We wondered how well these results correlate with our transcriptional data, given the differences in approaches. For the most part, our results agree, with the protein set comprising a subset of the genes identified in each tissue (Figure 2B); relatively few proteins (0.3% – 4.2%) are consistently detected only in the proteomic samples.

### Gene subset expression in adult tissues

To start to build models of activity within and communication between tissues, we examined the expression of particular gene classes (Figure 2C). The expression of transcription factors, including nuclear hormone receptors, is evenly distributed across tissues, while neurons have the highest counts of serpentine receptors and insulins (Figure 2C, Table S7), perhaps unsurprising given their role in interpreting and communicating the environmental state to the rest of the animal. Neurons and intestine express a high number of transmembrane proteins, and the intestine also expresses the highest number of secreted proteins, suggesting roles in intercellular communication. Insulins, while expressed most highly in neurons, can be found in every tissue, including hypodermis (Figure 2C, D), consistent with previous reports (Pierce et al., 2001; Murphy et al., 2007; Ritter et al., 2013). The further characterization of the tissue specificity of these factors will be helpful in building models to describe how the worm’s adult cells communicate with one another.

### Super-enriched genes characterize individual tissues

We next identified the genes that were specific for each tissue, that is, highly enriched relative to all other tissues (defined as genes that are significantly differentially expressed relative to the average expression in other tissues (FDR ≤ 0.05, log_FC_ > 5)) (Table S8). By Spearman correlation analysis of super-enriched gene expression (Figure S1C), neurons are the most distinct tissue. The GO terms for neurons and muscle (Figure 3A, Table S2) are largely expected: muscle’s super-enriched genes are associated with *muscle contraction*, *actin filament-based process*, *cell migration*, *calcium ion homeostasis*, *actin cytoskeleton organization*, *integrin signaling*, etc., while the neuronal super-enriched set is associated with many well-characterized neuronal functions, including *learning or memory*, *synaptic transmission*, and *neurotransmitter transport* (Figure 3A, Table S2). GO analysis of the intestine suggested that in addition to expected categories associated with digestion (*peptidase activity*, *proteolysis*), response to bacteria (*defense response*, *response to biotic stimulus*), and lipid metabolism (*localization* and *storage*), terms associated with the molting cycle, driven by the high expression of several collagens, appeared in the intestine super-enriched gene list (Figure 3A, Table S2).

**Figure 3:**
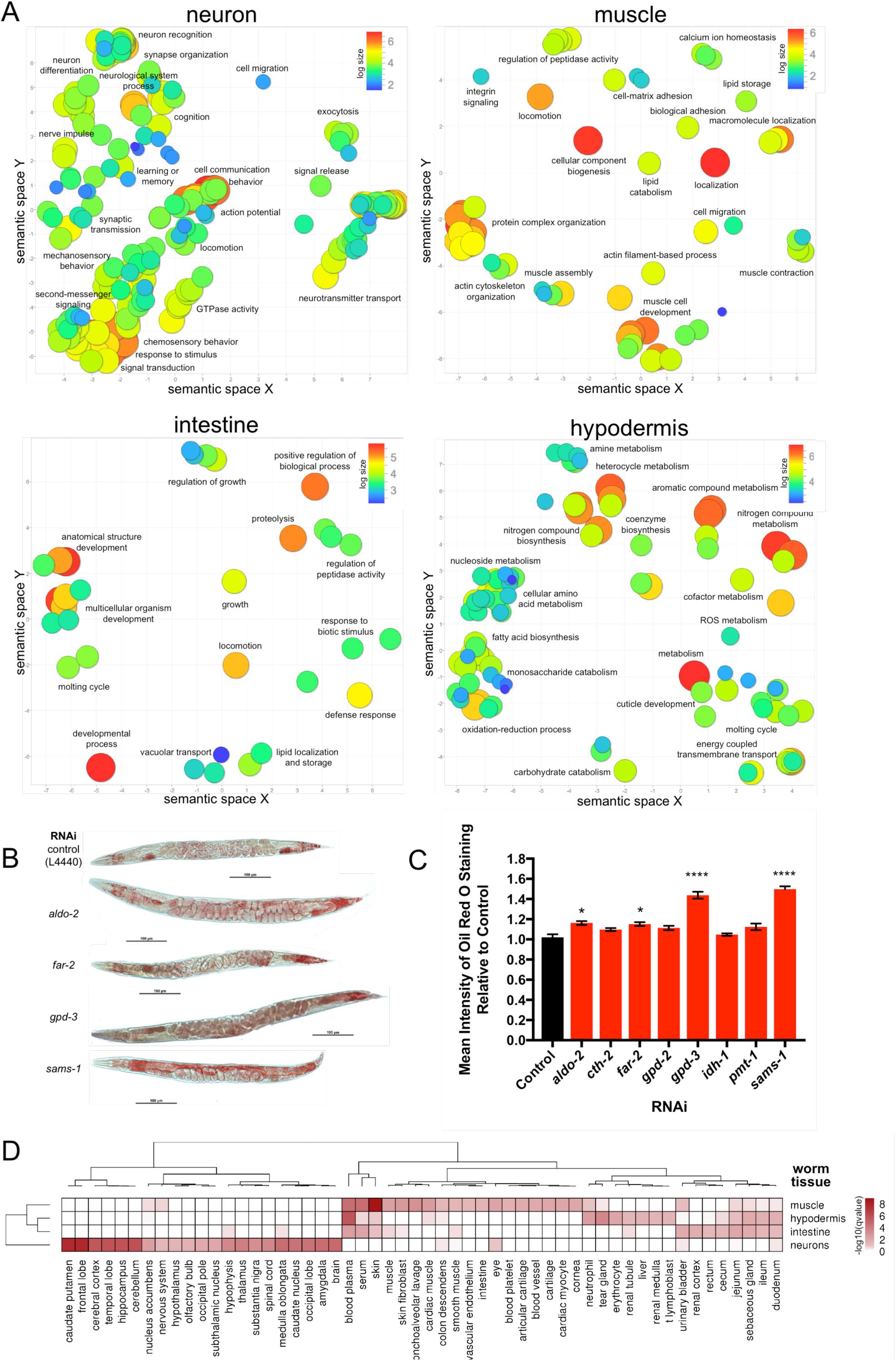
Comparisons and characterization of super-enriched gene sets. A) Gene ontology analysis of each super-enriched tissue list (Table S2). GO terms with padj < 0.05 (hypergeometric test) were plotted using REVIGO. B) Representative 20x images of Day 1 RNAi-treated animals after 6 hours of Oil Red O staining. No significant differences in worm size were observed. C) Mean intensity of Oil Red O staining analyzed using CellProfiler. *aldo-2*, *far-2*, *gpd-3*, and *sams-1* RNAi-treated animals had significantly more fat content (* p-value < 0.05, **** p-value < 0.0001 by one-way ANOVA), compared to vector L4440 control. D) Comparison of each worm tissue (human orthologs of muscle, hypodermis, intestine, and neuron superenriched gene lists) to human tissues (HPRD gene annotations (Keshava Prasad et al., 2009)). Human tissues with q-value < 0.05 (hypergeometric test) were plotted (values plotted are −log_10_(q-value), white = not significant, dark red = highly significant enrichment).

The *C. elegans* hypodermis is a relatively uncharacterized tissue, best understood for its role in collagen expression of the cuticle and molting during development (Cox et al., 1981), and those GO terms are represented here (Figure 3A, Table S2). However, it is interesting to note that the vast majority of GO terms associated with the hypodermal super-enriched genes are related to different types of metabolism, including *carbohydrate*, *amino acid*, *fatty acid*, *monosaccharide*, *nucleoside*, *ROS*, *co-factor*, *nitrogen compound*, *aromatic compound*, and *heterocycle metabolism*, as well as *co-enzyme biosynthesis* and *oxidation-reduction process*. These data suggest that the hypodermis, a syncytial tissue that makes up a substantial fraction of the adult animal’s body, may be a major site of metabolic processes in *C. elegans*. To test this hypothesis, we chose top-ranked hypodermal genes with predicted metabolic functions and available RNAi clones, knocked down their expression, and found that reduction of *aldo-2* (fructose-bisphosphate aldolase), *far-2* (fatty acid binding protein), *gpd-3* (glyceraldehyde 3-phosphate dehydrogenase), and *sams-1* (S-adenosyl methionine synthetase) significantly increased fat levels (Figure 3B-C, S2).

Motif analysis of the super-enriched genes for each tissue identified several candidate tissue-specific transcription factors (TFs) (Table S9). Hypodermal genes were enriched for binding sites of known hypodermal TFs, including ELT-3, BLMP-1, and NHR-25, and both the hypodermis and intestine super-enriched genes showed significant overrepresentation of TGATAAG, the DAF-16 Associated Element, which PQM-1 binds (Tepper et al., 2013); all four of these transcription factors are in our hypodermis expressed list (Table S1). The intestine was enriched for motifs associated with EGL-27 and ELT-7, while muscle was enriched for JUN-1, DMD-6, LIN-29, and ZTF-19/PAT-9. Neuron-enriched motifs include SKN-1/Nrf2, whose activity in neurons regulates longevity (Bishop and Guarente, 2007; Mizunuma et al., 2014), and MBR-1 (honeybee Mblk-1 related factor), which is expressed in the AIM interneuron, the site of long-term memory formation (Kage et al., 2005; Lakhina et al., 2015).

### Identification of orthologous worm/human tissues

*C. elegans* can be used as a model to study human disease due to its high degree of genetic, proteomic, cellular, behavioral, and tissue conservation. Functional comparisons are often used to draw parallels between worm and human tissues; for example, fat storage in the worm intestine and hypodermis is comparable to the function of human adipose tissue (Jones and Ashrafi, 2009), and the worm hypodermis, forming an outer protective layer covered in cuticle, is used as a model for human skin (Chisholm and Hsiao, 2012). Using our tissue-specific sequencing data, we compared transcriptional profile similarities of worm and human tissues. In an unbiased approach, we identified the human orthologs (Ruan et al., 2008) of genes in our super-enriched gene lists and compared them with curated tissue-specific gene annotations from the Human Protein Reference Database (Keshava Prasad et al., 2009) for significant overlap (hypergeometric test). Not surprisingly, worm neurons are transcriptionally most similar to various human brain regions and neuronal tissues, and worm muscle is similar to human muscle (Figure 3D, Table S10). Worm muscle also corresponded with human skin, sharing genes with GO terms related to basement membranes and cell migration, such as laminins, osteonectin, and integrins. Worm muscle and human skin also shared genes related to retinoic acid binding proteins (*lbp-1, -2, -3/* CRABP1 and 2) (Table S10). The worm intestine showed correlation with human small and large intestine, as well as parts of the human kidney, including the urinary bladder and renal cortex, sharing genes involved in the renin-angiotensin system (*acn-1*/ACE2, *asp-3*/REN) and various solute carrier family proteins. Surprisingly, the worm hypodermis exhibited no significant similarity to human skin, but rather correlated with liver and blood plasma, driven largely by metabolic genes (e.g., *ahcy-1*, *sams-1*, *cth-2*; Table S10). Hypodermis also correlated with immune cells, such as neutrophils and T cells, potentially related to the immune functions of the worm hypodermis (Figure 3D) (Taffoni and Pujol, 2015); the worm hypodermis and human neutrophils share genes involved in microbial responses such as *ZC416.6*/LTA4H (leukotriene A4 hydrolase), *sta-2*/STAT3, *adt-2*/CFP (ADAMTS metalloprotease/complement regulator), *ftn-2*/FTL (iron metabolism), and *ctl-2*/CAT (catalase) (Chen et al., 1994; Johnson and Wessling-Resnick, 2012; Melo and Ruvkun, 2012; Vigneshkumar et al., 2013; Zhang et al., 2015). These cross-species tissue similarities may be useful in developing new *C. elegans* models of human disease.

### Adults express tissue-specific isoforms

Previously, global tissue-specific isoform expression has been difficult to assess because most experiments have used whole worms, have detected gene expression using isoform-indiscriminant microarray platforms, require affinity purification (Gracida and Calarco, 2017), or only utilize short reads biased toward the 3’ ends of mRNA (Cao et al., 2017). Here we used relatively long reads with good coverage through most gene bodies (Figure 1C), and thus were able to assess the levels of expression of individual exons in each tissue. We identified 23,087 exons with significant tissue-specific alternative splicing (Table S11). Several well documented tissue-specific splicing events are corroborated by our data; for example, *unc-32*, a vacuolar proton-translocating ATPase, exhibits mutually exclusive exon splicing at two separate locations (exons 4 A/B/C and 7 A/B) (Kuroyanagi et al., 2013), and our data support the selective neuronal enrichment of exons 4B and 7A compared to other tissues (Figure 4A). Other examples include *egl-15’s* muscle-specific inclusion of exon 5A and repression of exon 5B (Kuroyanagi et al., 2006) (Figure 4B), and the ubiquitous expression of *unc-60* isoform A compared to the muscle-specific enrichment of isoform B (Ono et al., 2003) (Figure S1D).

**Figure 4:**
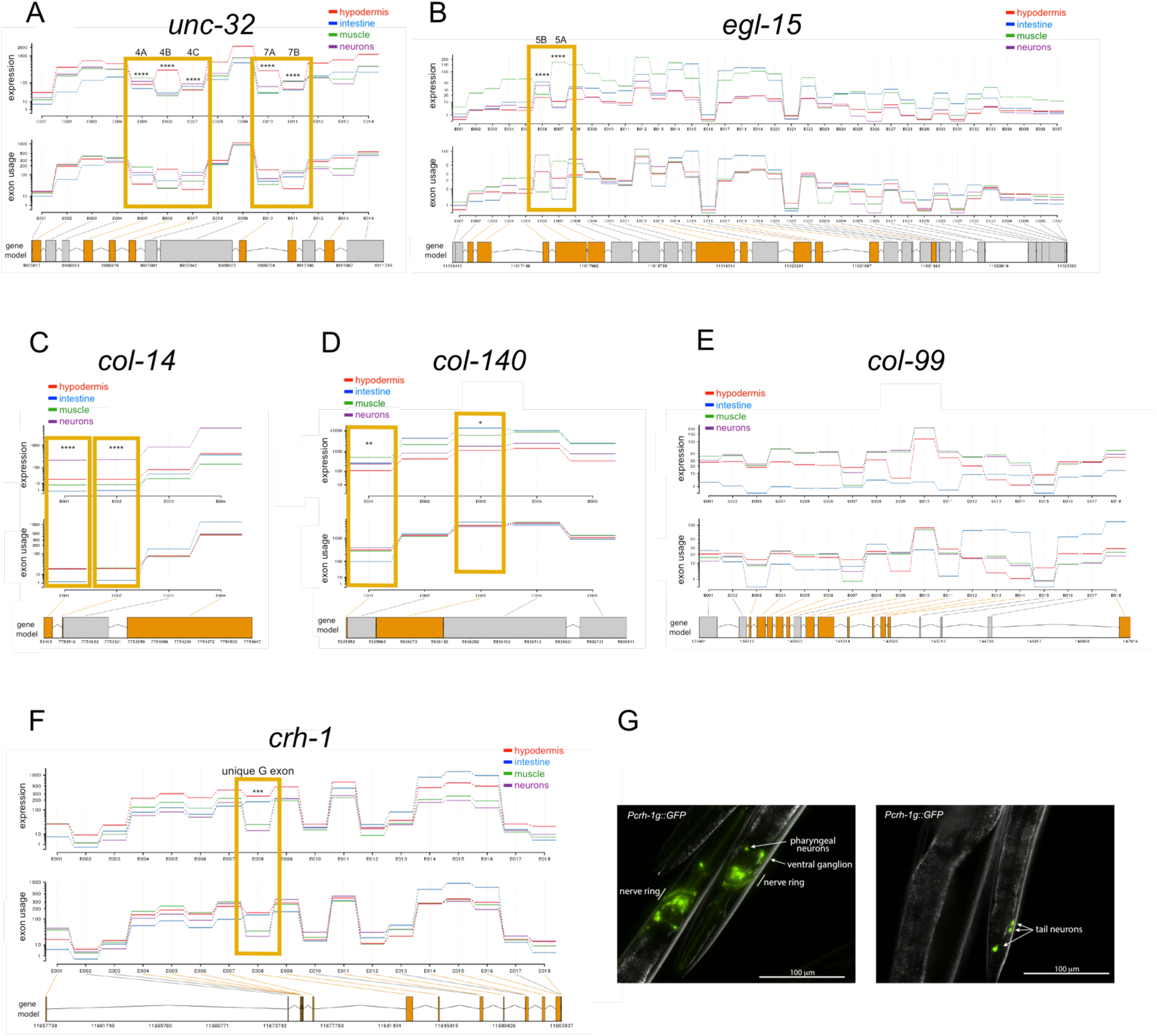
Alternative splicing analysis. DEXseq was used to identify significant splicing differences among tissues. Alternative splicing of *unc-32* (A) and *egl-15* (B) across tissues as examples that correlate well with previous reports. C-E) Collagens are expressed in multiple tissues, and some are spliced in a complex manner. F) Transcripts of a particular isoform of CREB, *creb-g*, are enriched in neurons. G) *Pcrh-1g::gfp* expression confirms that the *creb-g* isoform is expressed exclusively in several head and tail neurons. * p<0.05, ** p<0.01, *** p<0.001, **** p<0.0001.

We previously found that collagens play an unexpected but important role in the regulation of lifespan (Ewald et al., 2015). Here we found that a surprising number of collagens are alternatively spliced, with expression not just in the hypodermis as anticipated, but also in neurons, muscle, and intestine (e.g., *col-14;* Figure 4C). While most collagens have relatively simple exon structures, including those with differential splicing such as *col-140* (Figure 4D), some are complex; for example, *col-99* has at least 18 exons, with isoform expression in hypodermis, intestine, muscle, and neurons (Figure 4E). The role of tissue-specific collagen isoforms offers an interesting avenue for further study.

CREB, the cAMP response element binding protein, is an important and highly conserved transcriptional regulator of both metabolism and long-term memory (Bourtchuladze et al., 1994; Kauffman et al., 2010; Lakhina et al., 2015). CREB’s memory transcriptional targets are distinct from those involved in metabolism (Lakhina et al., 2015). Although *crh-1*/CREB is expressed throughout the worm, its activation specifically in the AIM interneuron is required for longterm memory (Kauffman et al., 2010; Lakhina et al., 2015). While the *crh-1a* isoform is expressed throughout the worm body (I. Hope, Wormbase), we found that *crh-1*/CREB has one isoform, *crh-1g*, that is highly enriched in neurons (Figure 4F). To test this model, we expressed GFP downstream of the promoter region adjacent to the unique 5’ *crh-1g* exon (*Pcrh-1g::gfp*), and found that unlike the *crh-1a* isoform, *crh-1g* is expressed exclusively in a subset of head and tail neurons, including neurons in the ventral ganglion where the longterm memory neuron AIM resides (Figure 4G).

### IIS//FOXO target expression in adult tissues

One of the applications of our tissue-specific transcriptional information is to identify tissue-specific targets of genetic pathways that affect both autonomous and non-autonomous gene expression. Insulin/IGF-1 signaling (IIS) is one of the major longevity regulation pathways, controlling the nuclear translocation and subsequent transcriptional activity of the DAF-16/FOXO transcription factor (Kenyon et al., 1993). IIS acts both cell autonomously and non-autonomously to regulate lifespan (Apfeld and Kenyon, 1998; Libina et al., 2003; Wolkow et al., 2000), highlighting the need to distinguish the two sets of regulators. The whole-worm downstream targets of DAF-16 have been identified and many have been characterized (Murphy et al., 2003; Tepper et al., 2013). While the tissue-specific expression of a few of these targets is known (Kaletsky et al., 2015; Zhang et al., 2013), tissue-specific information is not available for the entire set of somatic targets.

Utilizing the canonical ranked list of IIS/DAF-16 targets(Tepper et al., 2013) at a 5% FDR cutoff, we determined the site of expression of these Class I and II targets (*daf-2* up-and down-regulated, respectively). The largest subset (34%) of IIS/DAF-16 target genes are expressed in all tissues (e.g., *gpd-3*, *M60.4*), echoing the *adult determination of lifespan* category in the GO analysis of the ubiquitous gene set (Figure 1F). 63% of the Class I and II targets are expressed in the intestine, which has been previously identified as a site of DAF-16 regulation (Libina et al., 2003; Tepper et al., 2013); 10% are expressed exclusively in intestine (e.g., *cdr-2*, *pept-1*, *dod-17*; Figure 5A, B, Table S12). Fifty percent of the targets are expressed in the hypodermis, with about 13% expressed exclusively there (e.g., *dod-3*, *btb-16*, *drd-50*; 3%) or shared just in the intestine and hypodermis (e.g., *mtl-1*, *hacd-1*, *ZK6.11*; another 10%). These data support the role of the hypodermis as a major site of insulin signaling and DAF-16 activity (Zhang et al., 2013). The most highly expressed gene in hypodermis, the Notch ligand *osm-11*, is secreted by the hypodermis, and its knockdown leads to a DAF-16-dependent lifespan increase (Dresen et al., 2015). Another top Class I target, the mitochondrial protein *icl-1/gei-7*, is essential for desiccation tolerance (Erkut et al., 2016) and lifespan extension (Murphy et al., 2003) of *daf-2* animals. Hypodermal genes from this list can affect lipid storage in the intestine; knockdown of *pmt-1* leads to an increase in lipid droplet size and an increase in *fat-7* levels, as is observed in *daf-2* animals (Li et al., 2011).

**Figure 5:**
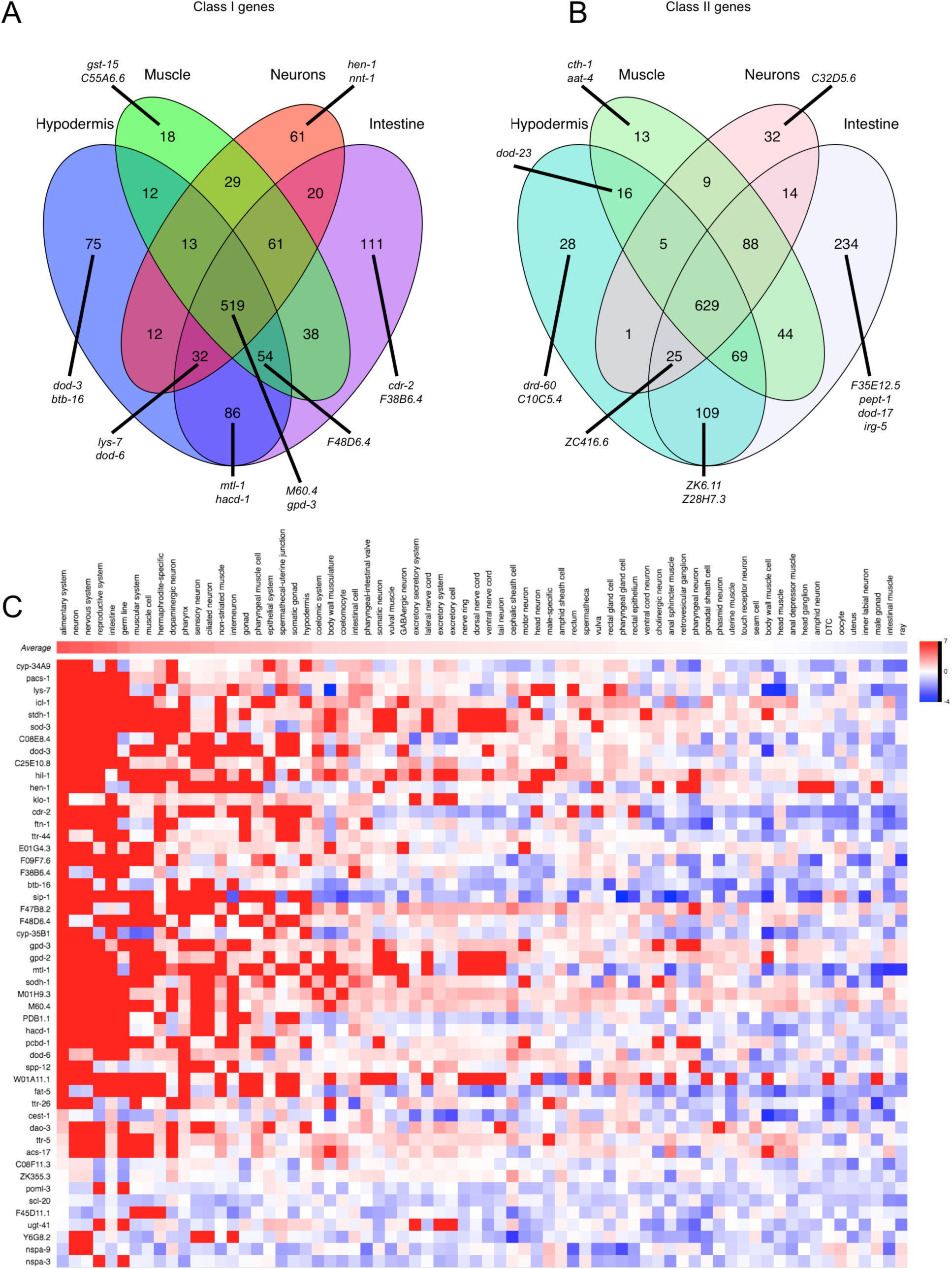
Tissue specificity of Insulin signaling/DAF-16 target expression. A) Class I (upregulated in long-lived *daf-2* worms, Tepper et al. 2013 at 5% FDR), and B) Class II (downregulated) tissue expression Venn diagrams show that 45-47% of previously identified targets are shared among all tissues. The highest fraction of Class I and II targets are expressed in the intestine and hypodermis. Examples of genes in each category, including the top 10 genes ranked in the previously published IIS/FOXO data (Tepper et al., 2013) are shown. C) The top 50 Class I genes (Tepper et al., 2013) were analyzed for predicted tissue expression (red = highest predicted expression, blue = lowest predicted expression). Gold standard annotations are represented with the highest score possible.

There are relatively few muscle-specific Class I or II DAF-16 targets (0.9%), and muscle subsets are all relatively sparse compared to the intestine and hypodermis. This may correlate with the observation that while DAF-16 expression in muscle rescues reproductive span (Luo et al., 2010), it does not rescue lifespan or dauer formation, unlike DAF-16 expression in intestine and neurons (Libina et al., 2003). Class I and II neuron-specific targets include *hen-1* and *nnt-1*, which are both highly regulated by the IIS pathway (Kaletsky et al., 2015; Murphy et al., 2003; Tepper et al., 2013). These findings are consistent with the documented and predicted (see below) expression patterns for Class I and Class II genes (Figure 5C, S8). Together, these data start to better delineate the tissue-specific functions of the IIS/DAF-16 pathway.

### Genome-wide expression predictions span 70 tissues and cell types

To create a more complete picture of spatio-temporal gene expression for tissues and life stages not yet characterized by direct sequencing analysis, we combined experimentally isolated tissue datasets with computational models of tissue-specific gene expression to generate genome-wide predictions across 70 diverse *C. elegans* cell types. This builds upon our previous machine learning approach that predicts gene expression across a few major tissues (Chikina et al., 2009), and is an advancement over other valuable but non-machine learning-based tools that rely solely on documented expression annotations (Angeles-Albores et al., 2016). We compiled 148,433 ‘gold standard’ gene annotations (Wormbase) and 4,372 publically available microarray and RNA-seq experiments across 273 datasets (including our tissue-ome dataset). To examine the quality of our predictions, we performed a hold-out analysis by using five-fold cross-validation, where a subset of known gene annotations is completely hidden from the algorithm as it is trained. Evaluating against the hidden annotations, we observed a median precision of 0. 8612 at 10% recall, corresponding to a median 6-fold improvement over the genomic background (Figure S3-6).

To further evaluate the adult tissue-ome (Figure 1), we generated predictions excluding the tissue-ome samples from our data compendium, then calculated average predicted tissue expression scores for each tissue’s super enriched genes (Table S2) across the 70 cell types. The predictions accurately characterized the identity of each sequenced tissue (Figure 6A). The predicted profiles in the 26 nervous system-related tissue expression models are especially striking, with very distinct, strong signals for the neuron super-enriched genes. Notably, the new tool outperformed our previous intestinal predictions (Chikina et al., 2009), while other large tissue predictions remained strong.

**Figure 6:**
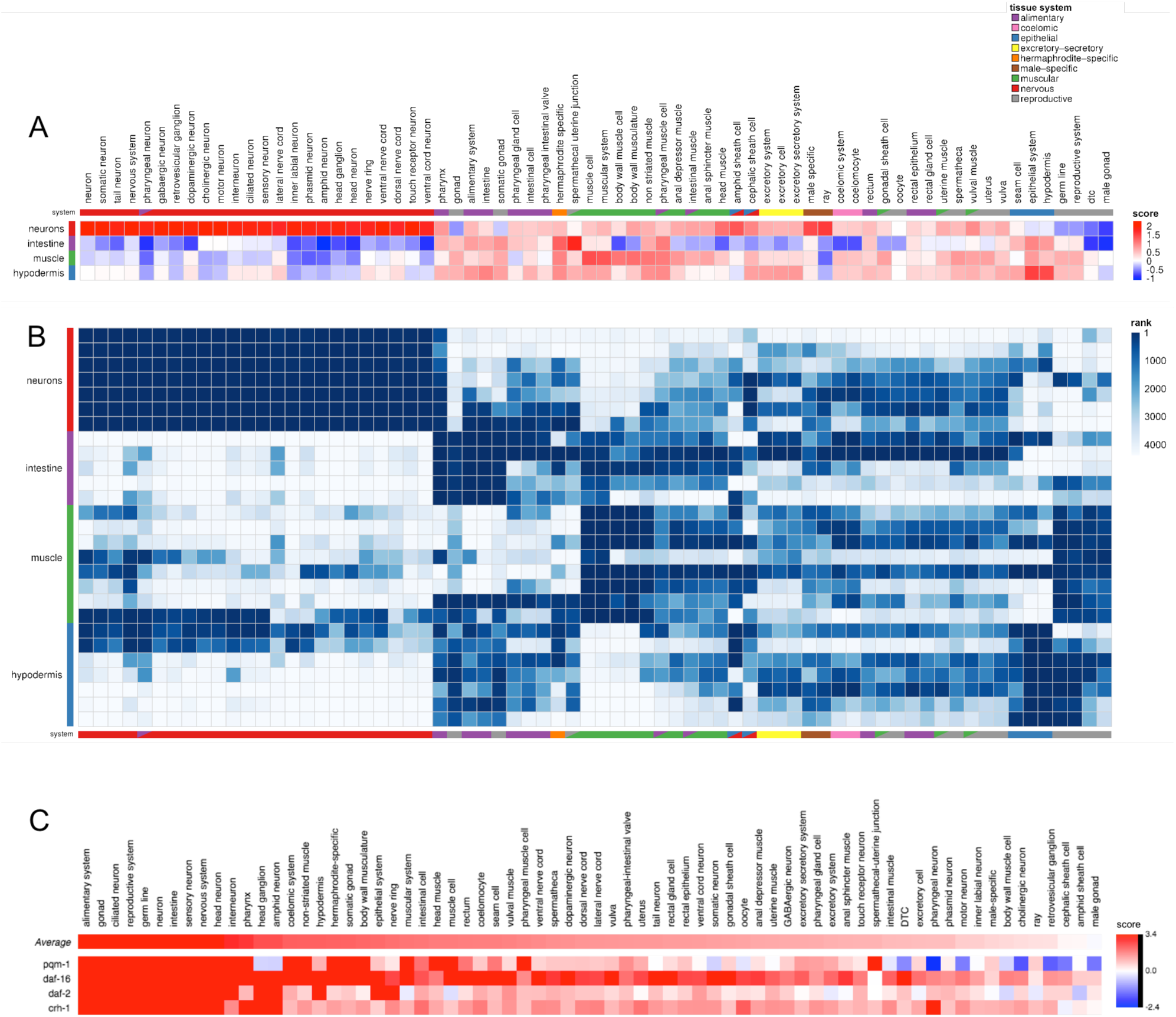
New prediction tool for sub-tissue/cellular gene expression. A) Concordance between predicted tissue expression scores and tissue-ome samples. Each row represents one of 4 tissue-ome tissues for which super-enriched genes were identified, and each column represents one of 70 tissues for which predictions were made. Columns were hierarchically clustered, and the tissue system(s) for each tissue is indicated. Each cell represents average predicted tissue expression (normalized SVM) score for corresponding super-enriched genes. B) Contribution of tissue-ome samples to tissue-gene prediction models. Each row represents one of 27 tissue-ome samples (across 4 tissues), and each column represents one of 70 tissues for which predictions were made. Row and column orderings are as in A. Each cell represents rank of sample in contribution to the corresponding tissue model. C) Tissue expression predictions for IIS-related genes (*daf-2*, *daf-16*, and *pqm-1*) and CREB (*crh-1*). Tissues are listed by average gene expression score.

We also explored the contribution of the tissue-ome to models based on the full data compendium. Taking advantage of the fact that for linear support vector machine (SVM) models, larger feature weights indicate higher relevance to the model (Fan et al., 2008), we examined the relative importance of each sample in the tissue-ome dataset. As expected, tissue samples were strongly upweighted in their relevant tissue models (Figure 6B) when compared to the other 4,314 samples in our data compendium.

### Applying the prediction tool

Individual and group gene expression predictions can be made using our interactive website (http://worm.princeton.edu). We examined several IIS pathway regulators to demonstrate the relevance of single gene predictions. Analysis of *daf-2* and *daf-16* predicts largely ubiquitous tissue distribution (Figure 6C), consistent with published data (Hunt-Newbury et al., 2007; Ogg et al., 1997) and the expression observed in the four major adult tissues (Table S1), as well as cell-autonomous functions of IIS in various tissues. The IIS/Class II gene-associated TF *pqm-1* (Tepper et al., 2013) exhibits broad expression, including in some neuronal subtypes and additional cells (Figure S8, Table S1), suggesting previously untested roles for this pathway.

To demonstrate the biological insight that can be gained from cell type expression prediction analysis of large gene sets, we analyzed published data sets of TGF-β, CREB, and DAF-16 signaling as examples. TGF-β Sma/Mab pathway signaling is highly conserved in worms and mammals, and regulates many diverse phenotypes including reproductive aging, body size, immune responses, and aversive olfactory learning (Luo et al., 2009; Mallo et al., 2002; Savage-Dunn et al., 2003; Zhang and Zhang, 2012). In *C. elegans*, the neuron-derived ligand DBL-1 modulates TGF-β transcription factor activity in the hypodermis, intestine, and pharynx. Hypodermal TGF-β signaling non-cell autonomously regulates oocyte quality underlying reproductive span (Luo et al., 2010), and analysis of TGF-β mutant (*sma-2*) oocytes revealed the suite of downstream targets important for maintenance of oocyte quality (Luo et al., 2010). Not surprisingly, these oocyte target genes are enriched for expression in oocytes and the reproductive system, as well as ubiquitously expressed genes (Figure S7A). By contrast, whole worm *sma-2* target genes from L4 animals lacking oocytes (Luo et al., 2010) are depleted for predicted reproductive system genes and are highly enriched in both predicted hypodermal and neuronal genes (Figure S7B). While whole-worm TGF-β pathway targets from L2 larval animals (Liang et al., 2007) share similar GO terms with L4 animals, such as collagens, metabolism, and hedgehog signaling (Luo et al., 2010), L2 TGF-β targets are not enriched for predicted neuronal genes (Figure S7C). This suggests that TGF-β transcriptional targets in late-larval animals, and perhaps adults, play a significant role in neuron function, which may explain the aberrant motor behaviors of *sma-2* and *sma-3* animals (Yemini et al., 2013), as well as the role of hypodermal TGF-β signaling in the neuronal processes regulating aversive olfactory learning (Zhang and Zhang, 2012). We also examined the transcriptional targets of CREB, whose activity in neurons is required for various forms of longterm memory (Bourtchuladze et al., 1994; Kauffman et al., 2010) and in metabolism (Altarejos and Montminy, 2011). The role of CREB in other tissues remains less well explored despite its broad tissue expression in *C. elegans* (Figure 6C, Table S1) and mammals, including humans (GTEx Analysis Release V7). We previously profiled Day 1 naïve and long-term memory-trained *crh-1/CREB* and wild-type animals to identify CrEB’s basal and memory target genes, respectively (Lakhina et al., 2015). CREB’s memory targets are highly enriched for genes with predicted neuronal expression (Figure 7A), and many of them are required for associative long-term memory (Lakhina et al., 2015). Interestingly, memory training also induced the CREB-dependent transcription of many genes strongly predicted to be selectively expressed in the hypodermis and intestine (Figure 7A); how these non-neuronal genes might influence memory is unknown. By comparison, a large cluster of CREB’s basal targets in young adult animals is enriched for expression in the reproductive system, with the highest average expression of targets predicted in the hypodermis, muscle, and intestine, but not neurons (Figure 7B). CREB is involved in mammalian spermatogenesis and oocyte maturation (Walker and Habener, 1996; Zhang et al., 2009), and CREB’s regulation of reproductive system expressed genes suggests that CREB may play a role in conserved pathways involved in reproduction and fertility.

**Figure 7.**
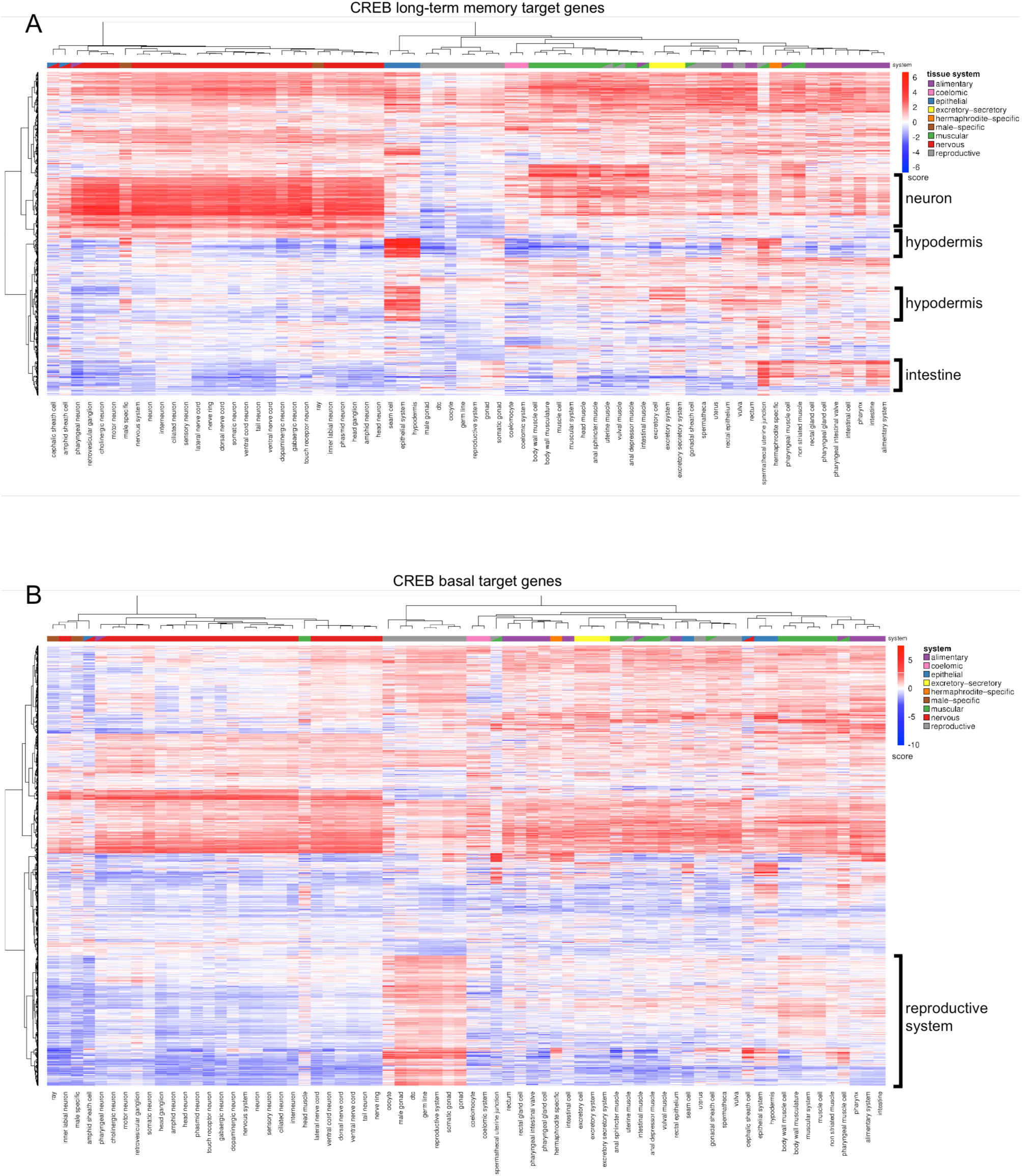
CREB-dependent long-term memory (LTAM) genes (A) and CREB-dependent naïve (basal) target genes (B) from Day 1 adult worms (Lakhina et al., 2015) were analyzed for predicted tissue expression. Prediction scores were used for cluster analysis using nonnegative matrix factorization (Gaujoux and Seoighe, 2010).

## Conclusions

*C. elegans’* simple architecture lends itself well to analysis of tissue-specific functions, and here we have exploited its simplicity to probe tissue-specific expression of adult animals. The isolation and RNA-seq analysis of *C. elegans* adults’ major somatic tissues has revealed that the transcriptomes of tissues differ from one another, and also from their counterparts in embryonic and larval stages, highlighting the importance of transcriptionally profiling specific developmental stages and cells relevant for the corresponding biology. These data provide the framework to explore the genetic basis of many phenotypes whose transcriptional underpinnings were not well represented with available larval data. The most striking example of this is the hypodermis, a tissue that is critically involved in development-specific larval molting processes, but in adults also becomes associated with metabolism, as shown here. These differences are likely to reveal the underlying basis of the hypodermis’ role in the regulation of reproductive aging, immunity, and longevity. The identification and prediction of insulin/FOXO target gene expression in non-intestine tissues, including hypodermis and neurons, lays the groundwork for understanding how lifespan signals are regulated within and across tissues. Moreover, the transcriptional conservation of the worm hypodermis and the human liver, in addition to other worm/human tissue similarities demonstrated here, provides new opportunities to model human diseases using genetic tools and resources of *C. elegans*.

We have provided tissue-specific transcriptional information, enrichment lists, GO analyses, and a new gene expression/cell type prediction tool for general usage. Our identification of tissue-specific isoform expression will enrich our understanding of differential uses for specific variants in different cellular contexts, and our prediction tool can be used to generate new hypotheses for gene function in specific cells. Together, enhance the ability to understand adult biological these methods and these gene expression data will function.

## Methods

### Strains and worm cultivation

OH441: otIs45[*Punc-119::GFP*], CQ163: wqEx34[*Pmyo-3::mCherry*], CQ171: [*Py37a1b.5::GFP*], BC12890: [*dpy-5(e907)I; sIs11337*(*rCesY37A1B.5::GFP* + pCeh361), SJ4144: *zcIs18* (*Pges-1::GFP*), CQ236: *Pcrh-1g::GFP* + *Pmyo-2::mcherry*. Worm strains were maintained at 20°C on HGM plates using *E. coli* OP50. Strains were synchronized using hypochlorite treatment prior cell isolation and grown to day 1 of adulthood on HGM plates with *E. coli* OP50.

### Adult cell isolation

Synchronized day 1 adult worms with GFP-labeled neurons, muscle, hypodermis, and intestine (*Punc119::GFP*, *Pmyo-3::mCherry*, *pY37A1B.5::GFP*, and *Pges-1::GFP*) were prepared for cell isolation, as previously described (Kaletsky et al., 2015).

### FACS isolation of dissociated cells

Neuron cell suspensions were passed over a 5 μm syringe filter (Millipore). Muscle and hypodermal samples were gently passed over a 20 mm nylon filter (Sefar Filtration). Intestinal cells were passed through a 35 mm filter and by spinning at 500 × g for 30s in a tabletop centrifuge. The filtered cells were diluted in PBS/2% FBS and sorted using a either a FACSVantage SE w/ DiVa (BD Biosciences; 488nm excitation, 530/30nm bandpass filter for GFP detection) or a BioRad S3 Cell Sorter (Bio-Rad; 488nm excitation). Gates for detection were set by comparison to N2 cell suspensions prepared on the same day from a population of worms synchronized alongside the experimental samples. Positive fluorescent events were sorted directly into Eppendorf tubes containing Trizol LS for subsequent RNA extraction. 30,000 to 200,000 positive fluorescent events were collected for each sample, depending on tissue. Both sorters were used for each tissue, and the type of sorter did not affect the distribution of samples by Principal Component Analysis (Figure 1B), suggesting that the sorter did not contribute to the variability between samples of a given tissue.

### RNA isolation, amplification, library preparation, and sequencing

RNA was isolated from FACS-sorted samples as previously described (Kaletsky et al., 2015). Briefly, RNA was extracted using standard Trizol/ chloroform/ isopropanol method, DNase digested, and cleaned using Qiagen RNEasy Minelute columns. Agilent Bioanalyzer RNA Pico chips were used to assess quality and quantity of isolated RNA. 10 to 100 ng of the isolated quality assessed RNA was then amplified using the Nugen Ovation RNAseq v2 kit, as per manufacturer suggested practices. The resultant cDNA was then sheared to an average size of ~200 bp using Covaris E220. Libraries were prepared using either Nugen Encore NGS Library System or the Illumina TruSeq DNA Sample Prep, 1 μg of amplified cDNA was used as input. RNA from a subset of samples was amplified using the SMARTer Stranded Total RNA kit-pico input mammalian, as per manufacturer suggested practices. No differences were observed between the two methods, and samples amplified by different methods clustered well (Figure 1B). The resultant sequencing libraries were then submitted for sequencing on the Illumina HiSeq 2000 platform. Sequences are deposited at NCBI BioProject PRJNA400796.

### Construction of promoter::gfp strains

The CREB (*crh-1*) isoform G promoter was made by cloning 1kb upstream of the *crh-1g* translation start site into the pPD95.75 Fire vector. N2 worms were injected with 25 ng/ul of pPD95-75-*Pcrh-1g::gfp* and 1 ng/ul of *Pmyo-2::mcherry* co-injection marker.

### RNA-seq data analysis

FASTQC was used to inspect the quality scores of the raw sequence data, and to look for biases. The first 10 bases of each read were trimmed before adapter trimming, followed by trimming the 3’ end of reads to remove the universal Illumina adapter and imposing a base quality score cutoff of 28 using Cutadapt v1.6. The trimmed reads were mapped to the *C. elegans* genome (Ensemble 84/WormBase 235) using STAR (Dobin et al., 2013) with Ensembl gene model annotations (using default parameters). Count matrices were generated for the number of reads overlapping with the gene body of protein coding genes using featureCounts (Liao et al., 2014). The per-gene count matrices were subjected to an expression detection threshold of 1 count per million reads per gene in at least 5 samples. EdgeR (Robinson et al., 2010) was used for differential expression analysis, and genes at FDR=0.05 were considered significantly differentially expressed. DEXSeq (Anders et al., 2012) was used for differential exon usage (splicing) analysis.

### Downsampling analysis

Count matrices of the aligned sequencing data were down-sampled using subSeq (Robinson and Storey, 2014). Reads were down-sampled at proportions using 10^∧^x, starting at x=−5 and increasing at 0.25 increments to 0. The down-sampled count matrices were used to assess stability of number of expressed genes detected at multiple depths (Figure S1A). Because of minimum library sizes for tractable differential exon usage analysis, reads with down-sampled proportions using 10^∧^x, from x=−2, increasing at 0.25 increments to 0 were used for assessment of power in detecting differential splicing (Figure S1B).

### Gene Ontology Analysis

Hypergeometric tests of Gene Ontology terms were performed on super-enriched gene lists; GO terms reported are a significance of *q*-value < 0.05 unless otherwise noted. REVIGO was used to cluster and plot GO terms with q-value < 0.05.

### Motif Analysis

RSAtools (Medina-Rivera et al., 2015) was used to identify the −1000 to −1 promoter regions of the super enriched genes and perform motif analysis. Matrices identified from RSAtools were analyzed using footprintDB (Sebastian and Contreras-Moreira, 2014) to identify transcription factors predicted to bind to similar DNA motifs.

### Oil Red O staining and analysis

Hypodermal genes appearing in metabolic GO terms were selected from the top of the super-enriched list (*aldo-2*, *gpd-2*, *sams-1*, *cth-2*, *pmt-1*, *idh-1*, and *fat-2*) or the expressed list (*far-2* and *gpd-3*) and knocked down using RNAi and compared to a vector (L4440) control. On day 1 of adulthood, all worms were stained in Oil Red O for 6-24 hours and then imaged using a Nikon Eclipse Ti microscope at 20x magnification (O’Rourke et al., 2009). Images were analyzed for mean intensity in fat objects using CellProfiler (Wählby et al., 2012).

### Worm-Human tissue comparison

Human orthologs (Ruan et al., 2008) of genes in our super-enriched gene lists were compared with curated tissue-specific gene annotations from the Human Protein Reference Database (Keshava Prasad et al., 2009) for significant overlap (hypergeometric test).

### IIS/FOXO target expression in adult tissues

The expression level (expressed defined as log(rpkm) >2) for previously published IIS/FOXO targets (Tepper et al., 2013, cut-off 5% FDR) were identified for each tissue. Tissue overlaps were graphed in Venn diagrams using the Venn diagram package in R.

### Expression data compendium assembly

To construct these models, we needed a large data compendium and high quality examples of tissue expression. We assembled 273 *C. elegans* expression datasets, representing 4,343 microarray and RNA-seq samples, including our tissue-ome library. All other datasets were downloaded from the Gene Expression Omnibus (GEO) data repository, ArrayExpress Archive of Functional Expression Data, and WormBase. Samples from each dataset were processed together (duplicate samples were collapsed, background correction and missing value imputation were executed when appropriate). Within each dataset, gene expression values were normalized to the standard normal distribution. All datasets were then concatenated, and genes that were absent in only a subset of datasets received values of 0 (in datasets in which they were absent). The predictions that were used to analyze the tissue-ome dataset were generated using a data compendium that excluded the tissue-ome library.

### Tissue expression gold standard construction

Gene annotations to tissues and cell types were obtained from curated anatomy associations from WormBase (Harris et al., 2014) (WS259) and other small-scale expression analyses as curated by Chikina et al. (2009). Only annotations based on smaller scale experiments (e.g., single-gene GFP, *in situ* experiments) were considered for the gold standard, excluding annotations derived from SAGE, microarray, RNA-seq, etc. Annotations were mapped and propagated based on the WormBase *C. elegans* Cell and Anatomy Ontology, where a stringent cutoff was used for which tissues and cell types were retained (>50 direct annotations and > 150 propagated annotations).

We defined a “tissue-slim” based on system-level anatomy terms in the WormBase anatomy ontology (immediate children of “organ system” and “sex specific entity,” under “functional system”). For each of the 70 tissues that were retained, a tissue-gene expression gold standard was constructed in which genes annotated (directly or through propagation) to the tissue were considered as positives. Genes that were annotated to other tissues, but not in the same tissue system, were considered negatives. Genes were assigned as positive or negative examples of tissue expression while taking into account the tissue hierarchy represented in the Cell and Anatomy Ontology.

### An interactive web interface to explore tissue-gene expression models

Our tissue-gene expression predictions and similarity profiles have all been made accessible at a dynamic, interactive website, http://worm.princeton.edu. From this interface, users can explore the predicted expression patterns of their gene(s) of interest. To facilitate this exploration, we have designed an interactive heatmap visualization that allows users to view hierarchically clustered expression patterns or sort by any gene or tissue model of interest. In addition, we also provide suggestions of genes with similar tissue expression profiles, which users can immediately visualize alongside their original query. All predictions and visualizations are available for direct download.

### Prediction and evaluation of tissue-gene expression profile and similarity

For each of the 70 tissues and cell types, we used the expression data compendium and corresponding gold standard as input into a linear support vector machine (SVM) to make predictions for every gene represented in our data. Specifically, given the vector of gene expression data (***x**_i_*) and training label (*y_i_*Î{−1,1}) for gene *i*, hyperplanes described by `w` and *b*, and constant *c*, the SVM’s objective function is: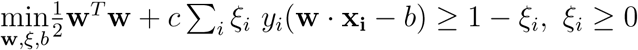, subject to the constraints: *y_i_*(w · x_**i**_ − *b*) ≥ 1 − *ξ_i_*, *ξ_i_* ≥ 0.

SVM parameters were optimized for precision at 10% recall under 5-fold cross validation. Resulting SVM scores were normalized to the standard normal distribution for any comparisons across tissues. Feature weights for each of the tissue SVM models were also retained for ranking and analysis of samples.

After applying the evaluated approach that we previously developed using a smaller set of data and tissues (Chikina et al., 2009), we used five-fold cross-validation to examine the quality of our predictions. Across all our tissues, we observed a median precision of 0.8612 at 10% recall, corresponding to a median 6-fold improvement over the genomic background (Figure S2, S3). Most of the neuron sub-types (e.g., cholinergic, dopaminergic, GABAergic, phasmid, amphid, ciliated, sensory, head, interneuron neurons, touch receptor neurons), as well as major tissues and tissue systems (e.g., alimentary system, reproductive system, nervous system) perform very well. Furthermore, in many of the small, less well characterized tissues, where there were fewer than 10% positives in our gold standard, we were able to substantially improve over the genomic background.

The tissue-ome dataset provides an opportunity to evaluate the predicted tissue-gene expression profiles on a snapshot of expression in neurons, hypodermis, intestine, and muscle. For the sake of comparing the predictions to the tissue-ome dataset, we also generated predictions while excluding the tissue-ome samples from our data compendium. We then calculated average predicted tissue expression scores based on each of these 70 tissue expression profile models across all identified super enriched genes.

## Acknowledgments

We thank the Murphy lab for valuable discussion, and the Center for *C. elegans* Genetics (CGC) for strains. CTM is the Director of the Glenn Center for Aging Research at Princeton and an HHMI-Simons Faculty Scholar. OGT is a senior fellow of the Genetic Networks program of the Canadian Institute for Advanced Research (CIFAR). This work was supported by the NIH (DP1 Pioneer Award (GM119167) and Cognitive Aging R01 (AG034446) to CTM, and R01 GM071966 to OGT), as well as by the Glenn Medical Foundation. VY and AR were supported in part by NIH T32 HG003284 and T32GM007388 grants. An early version of a portion of this work appeared in the doctoral thesis of A. Williams.

## Author Contributions

Conceptualization, R.K., A.W., V.Y., O.G.T., and C.T.M; Methodology, R.K., V.Y., O.G.T., and C.T.M; Investigation, R.K., V.Y., A.W. A.M.R., and S.B.K.; Writing, C.T.M and R.K. Funding Acquisition, O.G.T and C.T.M; Resources, O.G.T and C.T.M; Supervision, O.G.T. and C.T.M.

